# Preclinical evaluation of PHH-1V vaccine candidate against SARS-CoV-2 in non-human primates

**DOI:** 10.1101/2022.12.13.520255

**Authors:** Antoni Prenafeta, Gregori Bech-Sàbat, Alexandra Moros, Antonio Barreiro, Alex Fernández, Manuel Cañete, Mercè Roca, Luis González-González, Carme Garriga, Joachim Confais, Marion Toussenot, Hugues Contamin, Andrés Pizzorno, Manuel Rosa-Calatrava, Edwards Pradenas, Silvia Marfil, Julià Blanco, Paula Cebollada Rica, Marta Sisteré-Oró, Andreas Meyerhans, Cristina Lorca, Joaquim Segalés, Teresa Prat, Ricard March, Laura Ferrer

**Author notes:** These authors contributed equally (Antoni Prenafeta, Gregori Bech-Sàbat).

## Abstract

SARS-CoV-2 emerged in December 2019 and quickly spread worldwide, continuously striking with an unpredictable evolution. Despite the success in vaccine production and mass vaccination programmes, the situation is not still completely controlled, and therefore accessible second-generation vaccines are required to mitigate the pandemic. We previously developed an adjuvanted vaccine candidate coded PHH-1V, based on a heterodimer fusion protein comprising the RBD domain of two SARS-CoV-2 variants. Here, we report data on the efficacy, safety, and immunogenicity of PHH-1V in cynomolgus macaques. PHH-1V prime-boost vaccination induces high levels of RBD-specific IgG binding and neutralising antibodies against several SARS-CoV-2 variants, as well as a balanced Th1/Th2 cellular immune response. Remarkably, PHH-1V vaccination prevents SARS-CoV-2 replication in the lower respiratory tract and significantly reduces viral load in the upper respiratory tract after an experimental infection. These results highlight the potential use of the PHH-1V vaccine in humans, currently undergoing Phase III clinical trials.

## INTRODUCTION

Coronavirus disease 2019 (COVID-19), caused by severe acute respiratory syndrome coronavirus type 2 (SARS-CoV-2), emerged in Wuhan (China) in December 2019 and quickly spread worldwide^1^. COVID-19 has had a catastrophic economic and health impact worldwide despite governmental efforts to face the pandemic through preventive measures and rapid development of effective vaccines^2^. As of 14^th^ September 2022, over 605 million confirmed cases and over 6.4 million deaths have been reported globally^3^. Several vaccines have been approved for human use, including mRNA vaccines (BNT162b2, mRNA-1273), adenovirus vector-based vaccines (ChAdOx1 nCoV-19, Ad26.COV2.S, Ad5-nCoV-S, Gam-COVID-Vac), inactivated virus vaccines (VLA2001, PiCoVacc, BBIBP-CorV), and protein subunit vaccines (NVX-CoV2373, VAT00002)^4, 5^. The approved vaccines have shown a good safety and efficacy profile and their extensive use has resulted in a dramatic reduction in hospitalisations and mortality^6, 7^.

Nevertheless, the distribution of vaccines around the world is not equitable, leaving many low-and middle-income countries with an insufficient supply^8^. While 67.9% of the world population has received at least one dose of a COVID-19 vaccines, only 22.5% of people in low-income countries have received at least one dose^9^. Moreover, the emergence of new SARS-CoV-2 variants of concern (VOCs) with increasing immune evasion in vaccinated and convalescent individuals^10–12^, as well as the lack of vaccines that fully protect against reinfection and transmission^13^, has highlighted the need to develop new and effective SARS-CoV-2 vaccines. For instance, Omicron (BA.1) VOC contains many mutations in its spike protein associated with increased immune evasion and transmissibility ^14^. Indeed, two doses of ChAdOx1 nCoV-19 (AstraZeneca) or BNT162b2 (Pfizer-BioNTech) vaccine provided limited protection against infection with Omicron (BA.1) variant and mild disease^15^.

Second-generation vaccines are currently being developed to complement existing vaccines through heterologous prime-boost strategies and increased efficacy and availability. Subunit vaccines based on recombinant proteins can be produced on low-cost expression platforms and scaled easily at high yields, making them easier to produce and distribute globally. They are also stable and less reliant on a cold chain for their distribution than mRNA vaccines^16^, and they are associated with lower antibody-dependent enhancement (ADE) risks compared with other vaccine platforms^17^. The receptor-binding domain (RBD) of the spike protein is an attractive antigen for developing COVID 19 subunit vaccines^18^. RBD is the domain that binds to the human angiotensin converting enzyme 2 (ACE2) receptor to mediate viral entry. It is also the target for most of the neutralising antibodies elicited in natural infections, and several strong monoclonal antibodies have been identified in convalescent human sera and plasma ^19, 20^. Furthermore, most of the mutations of the Omicron (BA.1) spike protein are located at the RBD, including important mutations from previous variants^21^.

Our team has developed a ready-to-use adjuvanted recombinant RBD vaccine against SARS-CoV-2 referred to as “COVID-19 Vaccine HIPRA” or PHH-1V. The protein-based subunit vaccine candidate consists of a recombinant RBD fusion heterodimer of the Beta variant (B.1.351) and the Alpha variant of SARS-CoV-2 (B.1.1.7), produced in Chinese Hamster Ovary (CHO) cells and adjuvanted with an oil-in-water emulsion based on squalene (SQBA). Previous results showed that a prime-boost PHH-1V vaccination displayed an excellent safety profile in BALB/c and K18-hACE2 mice, produced RBD-specific antibodies with neutralising activity against Alpha, Beta, Delta and Omicron (BA.1) variants, and elicited a robust CD4^+^ and CD8^+^ T-cell response. Importantly, vaccination prevented mortality and body weight loss in SARS-CoV-2 infected K18-hACE2 mice, which is indicative of a high vaccine efficacy^22^. Currently, EMA’s human medicines committee (<u>CHMP</u>) has recommended authorising the PHH-1V vaccine (Bimervax) as a booster in people aged 16 years and above who have been vaccinated with an mRNA COVID-19 vaccine.

Non-human primates (NHPs) are often used to study viral vaccine efficacy and immunogenicity because most of innate and adaptive immune responses elicited against viruses are very similar to human responses^17^. Here, we report data on the immunogenicity, safety and efficacy profile of the PHH-1V vaccine candidate in twelve cynomolgus macaques (*Macaca fascicularis*) challenged with SARS-CoV-2 (D614G strain).

## RESULTS

### PHH-1V vaccination elicits high binding and neutralising antibody levels against SARS-CoV-2 in cynomolgus macaques

To assess the immunogenicity of PHH-1V vaccine, a group of six cynomolgus macaques (3 males and 3 females) were inoculated intramuscularly with 40 µg of the RBD fusion heterodimer mixed with SQBA on days (D) 0 and 21. A control group of same size was injected with PBS on the same dates. On D36, all animals were challenged intranasally and intratracheally with 2 x 10^6^ PFU of SARS-CoV-2 D614G strain and clinically monitored for 6 days. Sera from animals were obtained before each immunisation (D0 and D21), 7 days post-immunization (dpi) (D28), pre-challenge (D36) and 6 days post-challenge (dpc) (D42) to evaluate RBD-specific binding antibodies by ELISA (**Figure 1**).

**Figure 1.**
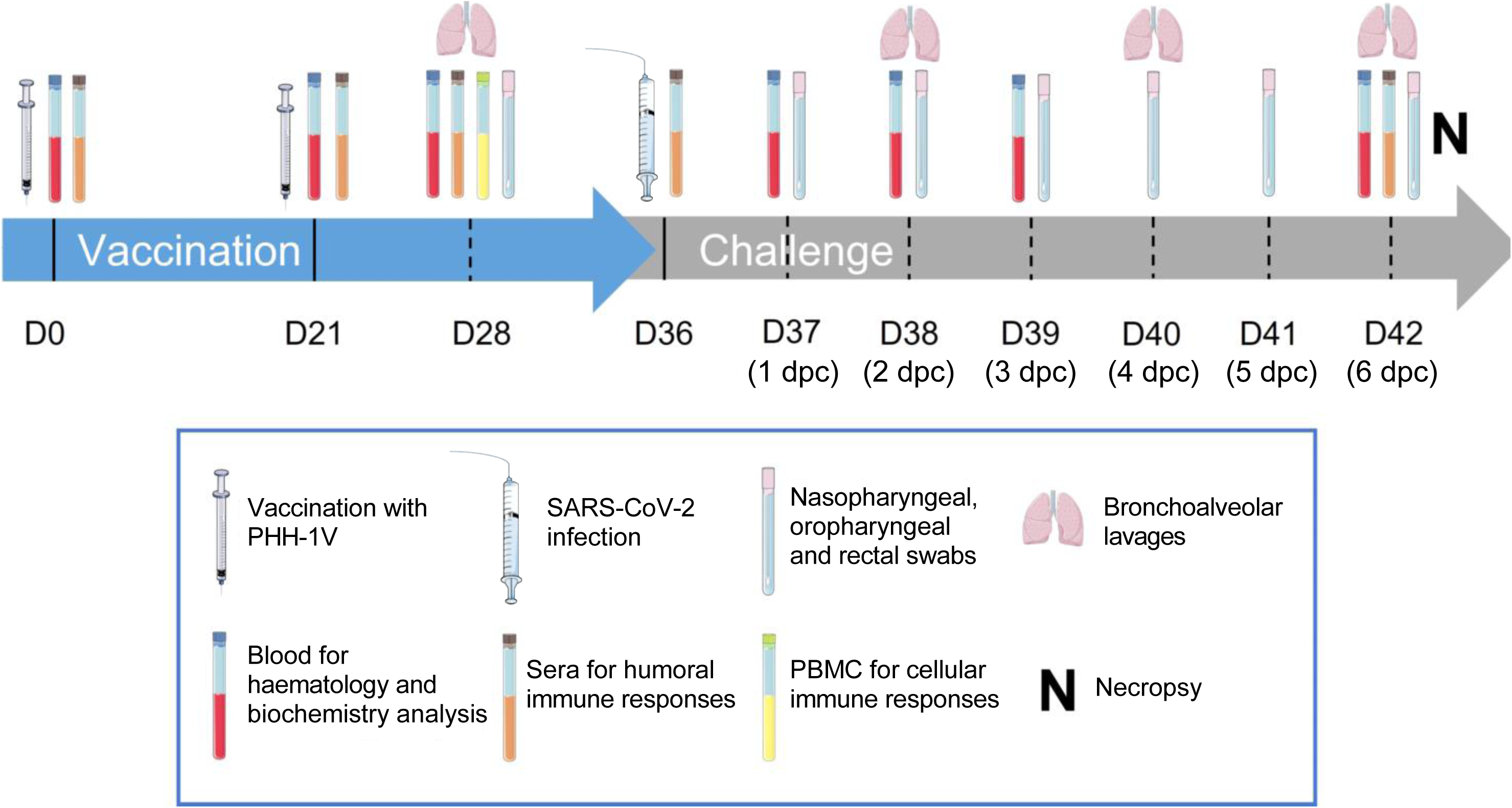
Study timeline. Cynomolgus macaques (6 per group) were immunised intramuscularly on days 0 and 21 with 40 µg of PHH-1V vaccine or with PBS as mock vaccine (control). On D36, animals were challenged intranasally and intratracheally with 2 x 10^6^ PFU of SARS-CoV-2. Different types of samples were taken at the timepoints indicated in the picture. All animals were euthanised on D42 (7 dpc). D, day; dpc, days post-challenge. Figure was generated using images assembled from Servier Medical Art (https://smart.servier.com).

Vaccination with one dose of PHH-1V induced significantly higher levels of IgG binding antibodies against RBD compared to control animals (p<0.01) on D21 (**Figure 2A**). A second dose of PHH-1V also significantly increased IgG antibody titres vs. control group (*p*<0.01) on D28 (7 dpi) and this superiority was maintained until the end of the study (D42) (**Figure 2A**). Additionally, IgA antibody titres against SARS-CoV-2 RBD were analysed in bronchoalveolar lavages (BAL) collected on D28 and D42 (6 dpc) (**Figure 2B**). Prime-boost immunisation with PHH-1V induced significant higher anti-RBD IgA antibody levels antibodies (p<0.05) on D42 compared to control groups. On D28, there is a tendency towards a higher anti-RBD IgA antibodies in PHH-1V-vaccinated animals.

**Figure 2.**
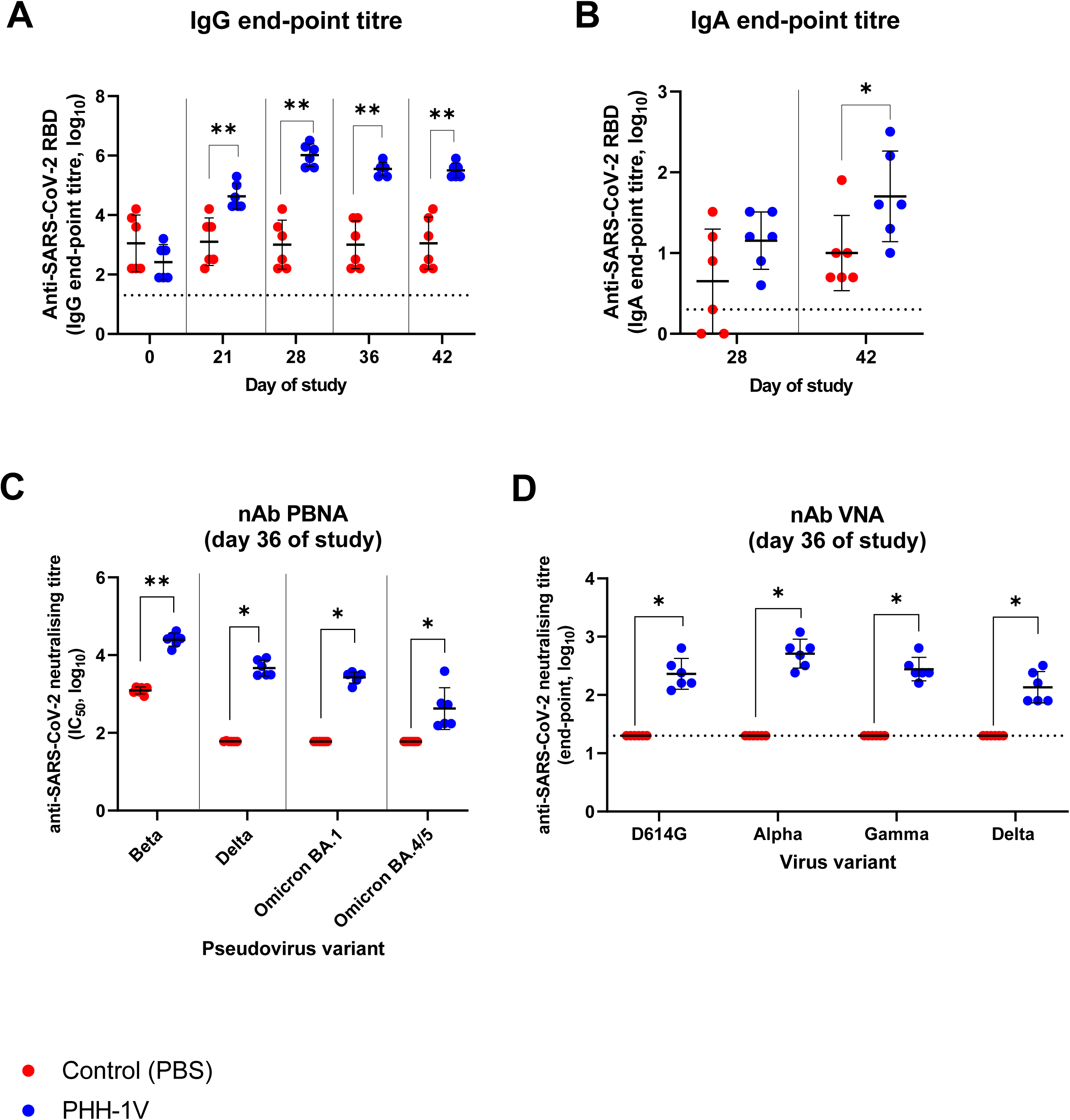
Humoral immune responses in cynomolgus macaques on PHH-1V vaccination. (A) IgG antibody titres against RBD were assessed by ELISA in sera from control animals (PBS, n=6) and vaccinated animals (PHH-1V, n=6) on days 0, 21, 28, 36 and 42. Log_10_-transformed data was analysed by means of a linear mixed effects model. (B) RBD-specific IgA antibody titres were assessed by ELISA in bronchoalveolar lavages (BAL) from control and PHH-1V vaccinated animals 7 days post-immunization (D28) and 6 days post-challenge (D42). Log_10_-transformed data were analysed using a Mann-Whitney’s U test. (C) Neutralising antibody response was analysed on D36 (pre-challenge) by pseudoviruses-based neutralisation assay (PBNA) against Beta, Delta, and Omicron (BA.1 and BA.4/BA.5) SARS-CoV-2 variants in macaques immunised with the PHH-1V vaccine (n=6) and control animals (PBS, n=6). Neutralising titres are expressed as log_10_ IC_50_, and limit of detection is shown as a horizontal dotted line. Each data point represents an individual animal with bars indicating the mean and the SD. Data were analysed using Mann-Whitney U-tests or one-sample Wilcoxon tests against the null H_0_: µ=1.80. (D) SARS-CoV-2 neutralising antibodies were analysed by the microneutralisation test (MNT) on D36 against D614G strain and Alpha, Gamma and Delta variants in macaques. Limit of detection is shown as a horizontal dotted line. Each data point represents an individual animal with bars indicating the mean and the SD. Data were analysed using one-sample Wilcoxon tests against the null H_0_: µ=1.30. (*p<0.05, **p<0.01).

SARS-CoV-2 neutralising antibody titres were determined initially in sera on D36 (pre-challenge) by Pseudovirus Based Neutralisation Assay (PBNA) using HIV reporter pseudovirus that express the spike protein of Beta, Delta and Omicron (BA.1 and BA.4/BA.5) VOCs (**Figure 2C**). PHH-1V vaccination induced higher neutralising antibody titres against Beta (p<0.01), Delta (p<0.05), Omicron (BA.1) (p<0.05) and Omicron (BA.4/BA.5) variants compared to control animals. Background neutralisation against the Beta variant was observed in non-vaccinated control animals, while no neutralising activity was observed in these groups against the Delta and Omicron variants. Neutralisation activity against SARS-CoV-2 D614G and VOC Alpha, Gamma and Delta was also confirmed on D36 by a microneutralisation test (MNT) with live viruses (**Figure 2D**). Animals vaccinated with PHH-1V showed a strong neutralising antibody response against all studied variants on D36 (p<0.05 vs. control group) as well as against SARS-CoV-2 D614G strain. No neutralising activity against SARS-CoV-2 and different VOCs was observed in animals inoculated with PBS.

### PHH-1V vaccination elicits robust T-cell immune responses against different SARS-CoV-2 VOCs in cynomolgus macaques

We next analysed specific T-cell immune responses induced by PHH-1V on D28 (7 dpi) in PBMCs stimulated with a RBD peptide pool from Alpha and Beta variants (**Figure 3A**) or Omicron (BA.1) variant (**Figure 3B**) through an ELISpot test. A prime-boost immunisation with PHH-1V induced significantly higher levels of specific IFNL- and IL-4-secreting cells against Alpha, Beta and Omicron (BA.1) variants on D28 compared to control group (p<0.01 and p<0.05 respectively for each stimulation). Remarkably, vaccinated animals showed a balanced Th1- and Th2-response against Alpha and Beta variants, but a polarised Th1-type immune response against Omicron (BA.1).

**Figure 3.**
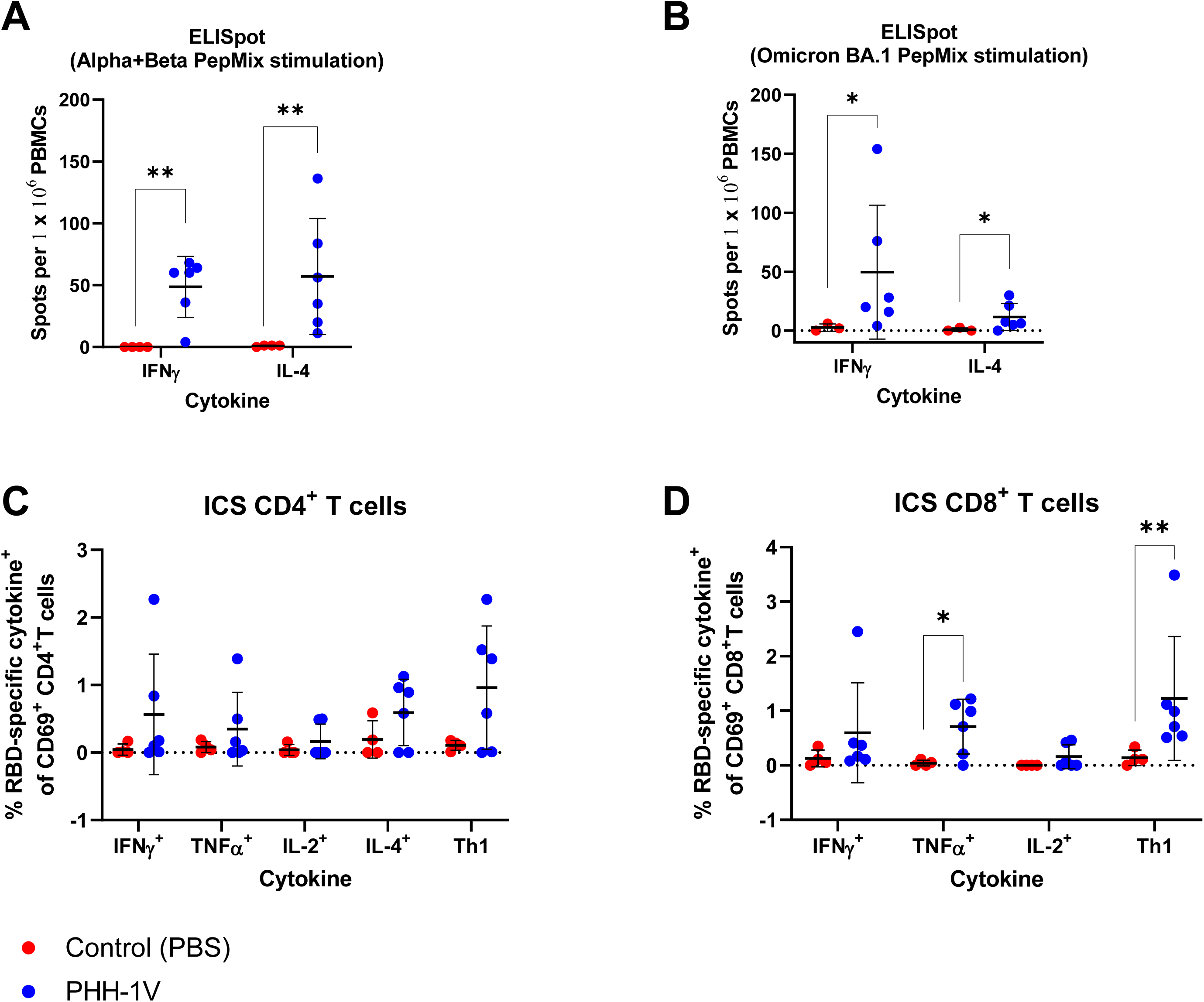
Cellular immune responses in cynomolgus macaques on PHH-1V vaccination. (A and B) PBMCs were isolated 7 days post-immunization (dpi) (D28) in PHH-1V vaccinated (n=6) and PBS-mock vaccinated control (n=3 or 4) group and then they were stimulated with an RBD peptide pool from Alpha and Beta variants (A) or Omicron (BA.1) variant (B) IFNL^+^-and IL-4^+^-expressing cells were determined by ELISpot assay. Arcsine-square root-transformed percentage data was analysed using generalised least squares models. (C and D) PBMC from vaccinated (PHH-1V, n=6) and control (PBS, n=4) cynomolgus macaques were isolated 7 dpi (D28). PBMCs were stimulated with an RBD peptide pool from Alpha and Beta variants and analysed by ICS. Frequencies of cytokine expressing CD4^+^ T cells (C) and CD8^+^ T cells (D) are shown. Frequencies of CD4^+^ and CD8^+^ T cells expressing Th1-like cytokines (IFNL, TNFα and/or IL-2) are also depicted. Arcsine-square root-transformed percentage data were analysed using generalised least squares models. Basal expression of cytokines stimulated with media was considered the background value and was subtracted from the peptide-specific response. Two animals from the control group were excluded in both assays (ELISpot and ICS) because one showed a consistent RBD-specific response and the other showed a higher percentage of dead cells in their PBMCs. In the ELISpot assay stimulated with Omicron (BA.1) peptides, another one was excluded for not having enough cells to perform the analysis. Each data point represents an individual animal with bars indicating the mean and the SD. Data were analysed using Welch’s permutation t-tests or Mann-Whitney U-tests. (*p<0.05, **p<0.01).

T-cell immune responses were further studied on D28 in PBMCs stimulated with a RBD peptide pool from Alpha and Beta variants by Intracellular Cytokine Staining (ICS). In PHH-1V-vaccinated animals, there was a trend toward higher specific CD4^+^ T-cells secreting Th1-like cytokines (IFNL, and/or TNFL, and/or IL-2) (**Figure 3C**). Animals vaccinated with PHH-1V elicited also robust and specific CD8^+^ T cells secreting TNFL (p<0.05) and Th1-like cytokines (IFNL, and/or TNFL, and/or IL-2; p<0.01) compared to animals inoculated with PBS (control) (**Figure 3D**). Likewise, IFNL^+^ CD4^+^ T-cell levels were similar to IL-4^+^ CD4^+^ T-cell levels, suggesting a balanced Th1- and Th2-response in animals vaccinated with PHH-1V (**Figures 3C**).

### PHH-1V vaccination protects from SARS-CoV-2 replication and infection in the upper and lower respiratory tract in cynomolgus macaques

To test PHH-1V efficacy on SARS-CoV-2 infection in cynomolgus macaques, we analysed the levels of genomic RNA (gRNA) to determine the viral load in nasopharyngeal, oropharyngeal and rectal swabs, as well as in bronchoalveolar lavage (BAL) by RT-qPCR (**Figure 4A-D**). PHH-1V vaccination significantly reduced viral gRNA levels in nasopharyngeal swabs 2 and 3 dpc (p<0.05) compared to PBS vaccination (**Figure 4B**). Furthermore, PHH-1V-vaccinated animals controlled SARS-CoV-2 infection in BAL and exhibited significantly lower gRNA levels 2, 4 and 6 dpc (p<0.05) compared to control animals (**Figure 4D**). At the end of the study, viral gRNA was not detected in BAL of three out of six vaccinated animals with PHH-1V, but could be detected in all six control animals.

**Figure 4.**
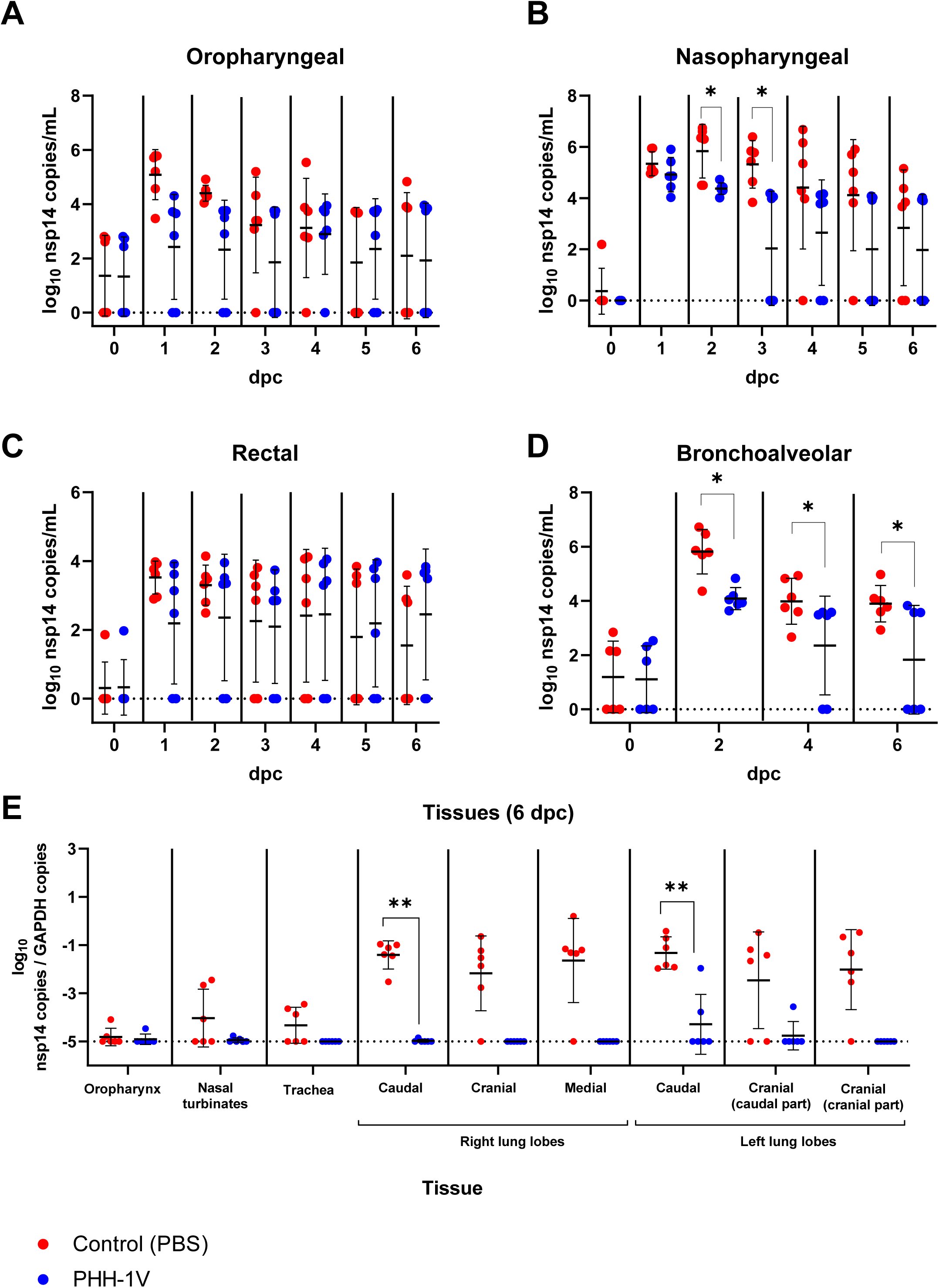
Quantification of viral RNA copies in lungs and upper respiratory tract of infected animals. (A-D) SARS-CoV-2 genomic RNA copies were measured by RT-qPCR in oropharyngeal swabs (A), nasopharyngeal swabs (B), rectal swabs (C) and BAL (D) from PHH-1V-vaccinated (n=6) and control (n=6) animals at different times post-challenge. Log_10_-transformed data were analysed using linear mixed effects models and gRNA is expressed as log_10_ (nsp14 copies/mL). (E) At 6 dpc (D42), animals were euthanised and viral RNA copies were measured in lungs, trachea, nasal turbinates, and oropharynx by RT-qPCR. Genomic RNA is expressed as log_10_ (nsp14 copies/GAPDH copies). Each data point represents an individual animal with bars indicating the mean and the SD. Data were analysed using Mann-Whitney U-tests or one-sample Wilcoxon tests against the null H_0_: µ = -5. (*p<0.05, **p<0.01). dpc, days post-challenge.

Six dpc all animals were euthanised, and some SARS-CoV-2 target tissues were collected to determine viral gRNA by RT-qPCR (**Figure 4E**). PHH-1V vaccinated animals exhibited significantly lower viral gRNA levels in lungs compared to PBS vaccinated animals, with a major decrease in the caudal lobes (p<0.01). Importantly, SARS-CoV-2 was undetectable 6 dpc in lungs, trachea and nasal turbinates in most animals immunised with PHH-1V vaccine.

We also analysed the subgenomic RNA (sgRNA) in nasopharyngeal swabs and BAL by RT-qPCR to confirm these results (**Figure S2**). While viral sgRNA was not detected in BAL from PHH-1V-vaccinated animals, it was detected in BAL from control animals 2 and 4 dpc (p<0.05). There was also a trend towards higher positive sgRNA samples in nasopharyngeal swab supernatants from control animals.

To further study PHH-1V efficacy in cynomolgus macaques, we also analysed the infective viral load of SARS-CoV-2 in nasopharyngeal and oropharyngeal swabs, BAL, and lungs by 50% tissue culture infectious dose (TCID_50_) assay. All PHH-1V-immunised animals controlled SARS-CoV-2 infection with no detection of infectious virus after challenge in oropharyngeal (**Figure 5A**) and nasopharyngeal swabs (except for one animal at 1 dpc, **Figure 5B**), BAL (except for one animal at 6 dpc, **Figure 5C**) and lungs (**Figure 5D**). In contrast, control animals exhibited high titres of SARS-CoV-2 in oropharyngeal up to 1 dpc, nasopharyngeal swabs and BAL up to 2 dpc and in lungs 6 dpc. Importantly, PHH-1V vaccination significantly reduced the infectious viral load in nasopharyngeal swabs 1 dpc (p<0.05) and in the cranial part of the cranial lobe (p<0.01) and caudal (p<0.05) lobes of lungs 6 dpc (**Figures 5B and 5D**).

**Figure 5.**
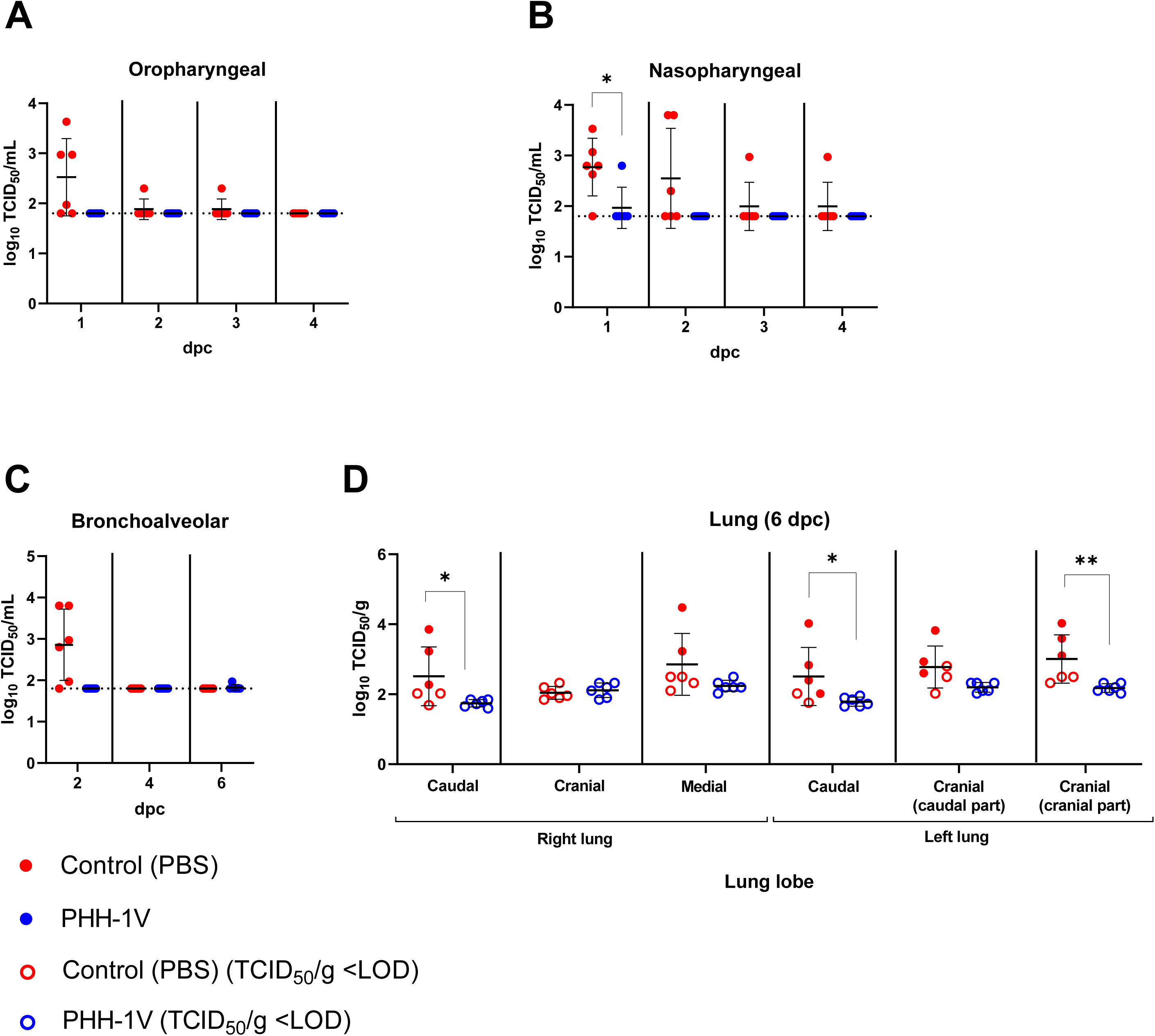
Quantification of infectious viral load in lungs and upper respiratory tract of infected animals. (A-C) Infectious viral load was measured by TCID_50_ assay in oropharyngeal swabs (A), nasopharyngeal swabs (B) and BAL (C) from PHH-1V-vaccinated (n=6) and control (n=6) animals at different times post-challenge. Results were expressed as log_10_ TCID_50_/ml, and the limit of detection (LOD) is indicated as a horizontal dotted line. Data were analysed using Mann-Whitney U-tests or one-sample Wilcoxon tests against the null H_0_: µ = 1.80. (D) At 6 dpc (D42), animals were euthanised and infectious virus were measured in lungs by TCID_50_ assay. Viral load is expressed as log_10_ TCID_50_/g. Hollow points indicate values under the limit of detection while solid points indicate values above the limit. Each data point represents an individual animal with bars indicating the mean and the SD. Data were analysed using Mann-Whitney U-tests. (*p<0.05, **p<0.01). dpc, days post-challenge.

### PHH-1V vaccination protects from lung disease induced by SARS-CoV-2 infection in cynomolgus macaques

Animal lungs were also processed after euthanasia 6 dpc in order to perform histopathological analysis. Lung inflammation scores of control animals were significantly higher (p<0.05) than PHH-1V vaccinated animals (**Figure 6A**). Likewise, the extent and severity of bronchointerstitial inflammation were more prominent in control animals (**Figure 6B, left**) than in those vaccinated with PHH-1V (**Figure 6B, right**), in which only mild and localised inflammation was detected. No findings were noted in histopathological examinations of the other analysed tissues, such as the trachea, oropharyngeal and nasal mucosa, lymph node, kidney, heart, liver and spleen (data not shown). Hence, the PHH-1V vaccine has a beneficial impact in reducing the lung disease caused by SARS-CoV-2.

**Figure 6.**
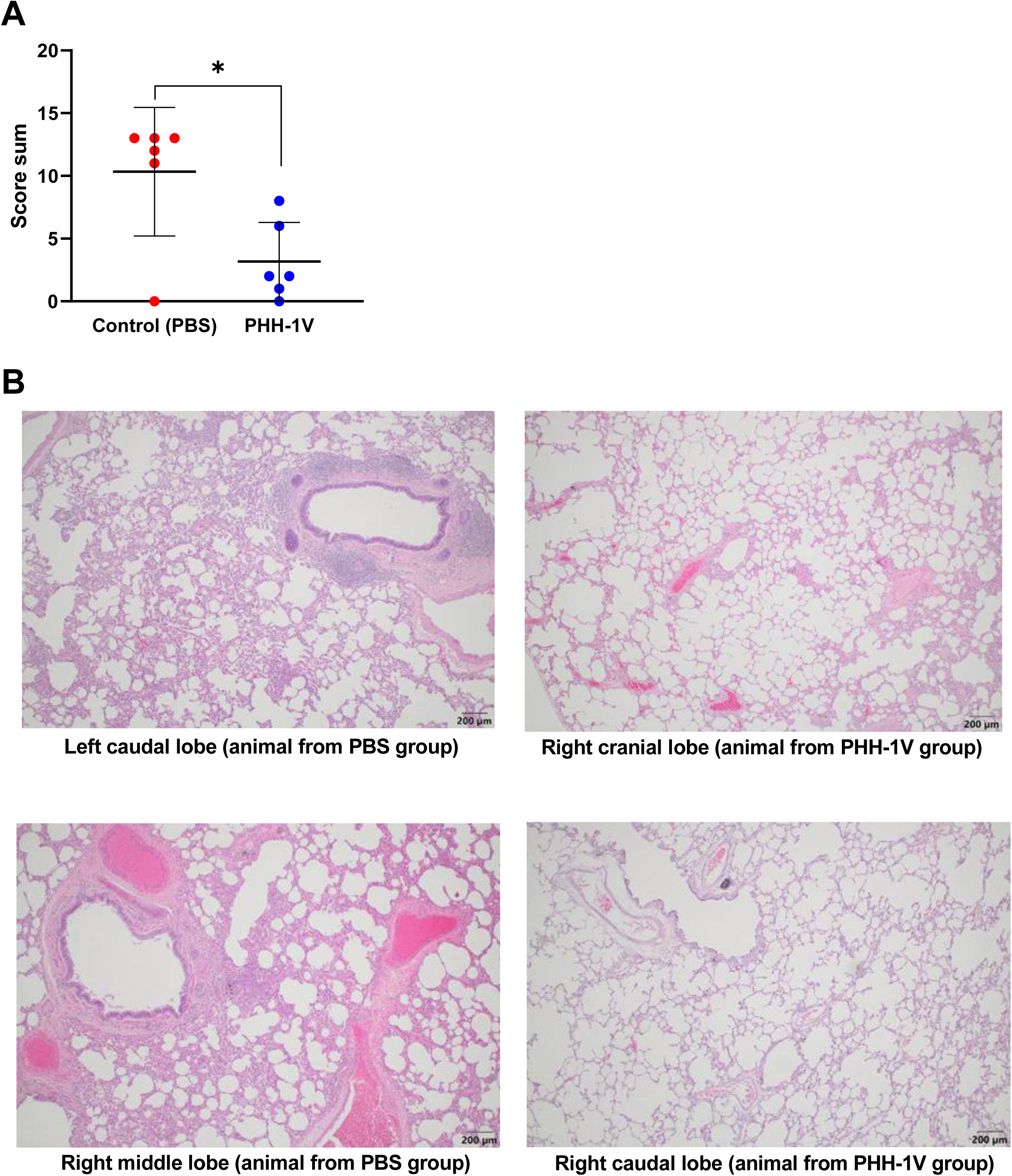
Lung histopathology in infected animals. Histopathological analyses were performed in lungs sections of 4 μm from PHH-1V-vaccinated and controls animals euthanised 6 dpi. (A) Total inflammation score calculated as the sum of 6 lobes (right caudal, left caudal, right cranial, left cranial [cranial part], left cranial [caudal part] and right middle). Each data point represents an individual animal with bars indicating the mean and the SD. Data were analysed using a permutation Welch’s t-test (*p<0.05). (B) Illustrative microphotographs of bronchointerstitial inflammation in one or more lung sections in males and females (200 µm). Lung sections from control animals on the left side, and sections from PHH-1V vaccinated animals on the right side.

Additionally, to confirm the SARS-CoV-2 replication in lungs, it has been performed an immunohistochemistry (IHC) technique to detect SARS-CoV-2 NP antigen in different lung sections from vaccinated and non-vaccinated animals (**Figure S3**). All non-vaccinated animals showing detectable virus by titration were positive by IHC in at least one lung section tested. In contrast, those animals immunized with PHH-1V that were analysed by IHC did not exhibit SARS-CoV-2 replication.

### Humoral and cellular immunity correlates with different efficacy parameters

To test the association between the immunogenicity and efficacy parameters, Kendall’s Rank correlation tests have been employed. The correlation between specific antibody and T cells levels and viral titre in nasopharyngeal and oropharyngeal swabs, BAL and different lung sections were assessed statistically by means of Kendall’s Tau-B (**Figure S4**). Overall, neutralising antibody titres against SARS-CoV-2 D614G (PBNA) and beta variant (VNA) inversely correlates with viral titres in upper respiratory tract (oropharyngeal swabs) and lower respiratory tract (BAL and lungs); and T cells against alpha, beta and BA.1 omicron variants inversely correlates with viral titres in upper respiratory tract (oropharyngeal swabs) and lower respiratory tract (BAL and lungs).

### PHH-1V vaccine is safe and well-tolerated in cynomolgus macaques

PHH-1V vaccine was administered intramuscularly. Local reactions at the injection site were monitored the day of the injection and daily up to 5 days after administration. No local reactions (oedema, induration, crust or haematoma) were observed at the injection site in any vaccinated animal, after either the first or second vaccination.

Animals were also monitored daily for clinical signs and twice daily for mortality from the first day of the acclimation phase to the end of the *in vivo* experimental phase. No clinical findings were recorded before or after the two administrations of either PHH-1V or PBS (control) on D0 and D21. No unexpected death or euthanasia for reaching endpoints occurred during the study in PBS and PHH-1V vaccinated animals.

Body temperature was monitored with a rectal thermometer around the vaccination and infection days on awake animals. Rectal temperatures were stable during the whole study before the SARS-CoV-2 experimental infection, with no differences between PBS and PHH-1V vaccinated animals. On D0 (prime vaccination) and D21 (boost vaccination), a lower temperature was recorded in the evening than in the morning in both groups (**Figure 7A**). This variation in body temperature is frequently observed in cynomolgus macaques and can be explained by the circadian rhythm^23^.

**Figure 7.**
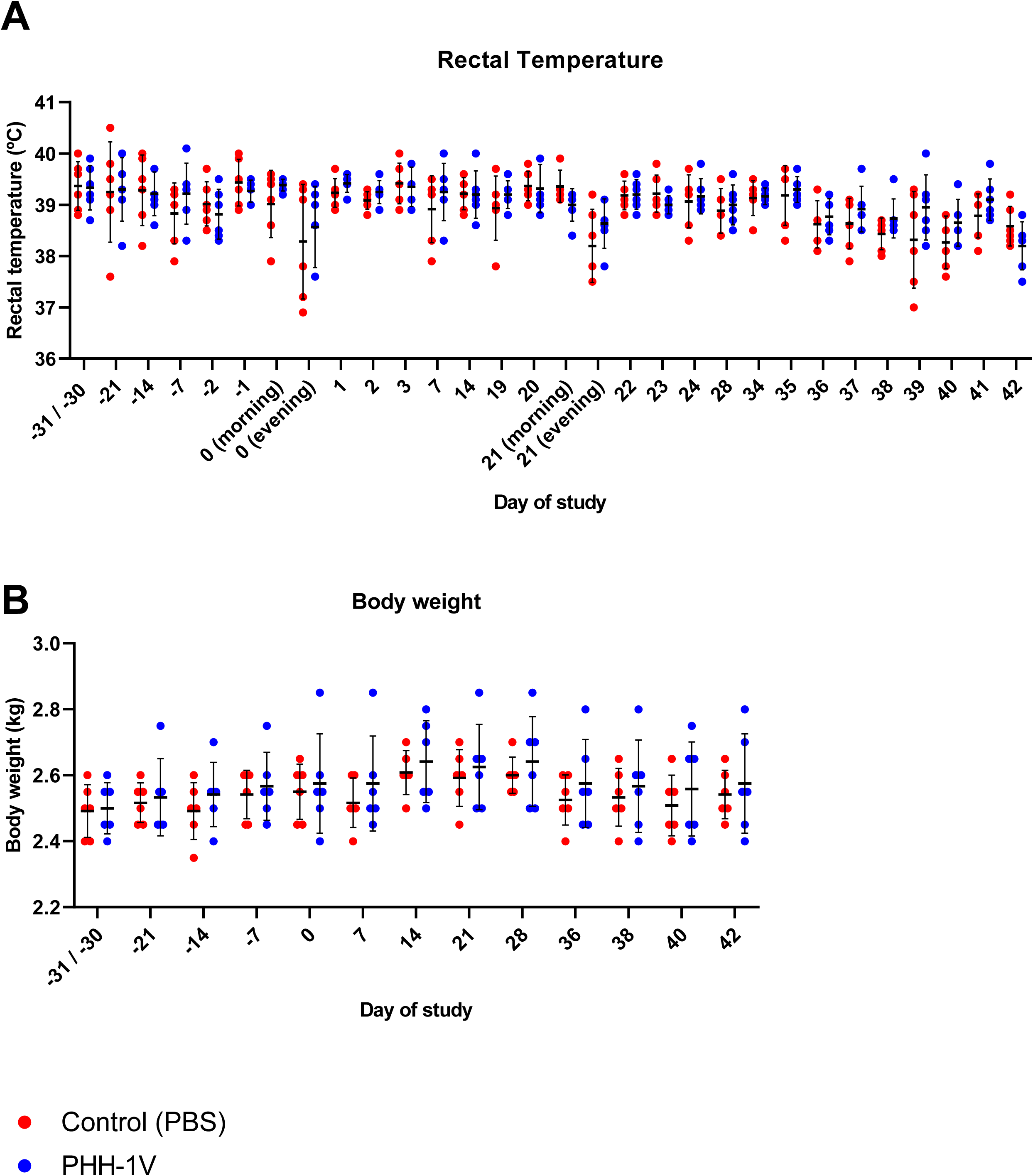
Safety assessment of cynomolgus macaques vaccinated with PHH-1V and PBS (control group). (A) Body temperature was monitored with a rectal thermometer around the administration and infection days in PHH-1V-vaccinated (n=6) and control (n=6) animals. Temperature data were analysed using a linear mixed effects model. (B) Animals were weighed once weekly throughout the acclimatization phase and during the vaccination phase. They were weighed three times a week after experimental infection. The figure shows body weight recorded throughout the study for each PHH-1V-vaccinated or control animal. Each data point represents an individual animal with bars indicating the mean and the SD. Body weight data were analysed using a linear mixed effects model.

Animals were weighed at least once weekly before SARS-CoV-2 infection, and three times a week after infection. Appetite and body weight were stable during the whole study before the infection, with no differences between PBS and PHH-1V vaccinated animals (**Figure 7B**). Nevertheless, consumption of food pellets during the week of infection was reduced in both groups. These observations are consistent with several experimental procedures performed that required frequent sedation and anaesthesia.

Finally, haematology and clinical chemistry levels were in the normal range (data not shown). Variations observed in some parameters are compatible with blood losses caused by multiple blood samples performed from D28. Vaccination with PHH-1V did not appear to have an impact on these parameters.

## DISCUSSION

To date, the scientific community and the pharmaceutical industry, with the support of both public and private sectors, have made an unprecedented global effort to develop and accelerate several vaccines worldwide to mitigate the COVID-19 pandemic. Despite the success of current vaccines, which have shown effectiveness in the prevention of severe disease, reduced infection rates and lowered mortality rates, the pandemic situation is not yet over. The emergence of new SARS-CoV-2 variants, characterised by enhanced immune evasion and transmissibility, together with the extremely unequal distribution of currently available vaccines, has made the virus difficult to contain^24^. This is of special importance in low- and middle-income countries, with only 22.5% of the population having received at least one vaccine dose^9^. Hence, it is critical to develop new second-generation vaccines effective against novel VOCs, which could be used as a booster to maintain and enhance immunity against SARS-CoV-2^25^.

Here, we have described the immunogenicity, efficacy, and safety of the PHH-1V vaccine candidate in cynomolgus macaques. PHH-1V vaccine is based on a recombinant heterodimer fusion protein comprising in tandem the RBD of the B.1.351 (Beta) and B.1.1.7 (Alpha) SARS-CoV-2 variants and formulated with an oil-in water emulsion based on squalene (SQBA). We have recently reported that PHH-1V vaccine elicited high humoral and cellular immune responses against several SARS-CoV-2 variants and was fully effective and safe in BALB/c and K18-hACE2 mice^26^. Most importantly, PHH-1V includes the sequences corresponding to two heterologous VOCs as part of the same protein molecule, which makes it differ from other adjuvanted protein-based subunit vaccines, such as NVX-CoV2373 (Novavax)^27^, ZF2001 (Anhui Zhifei Longcom Biopharmaceutical)^28^ or VAT00002 (Sanofi-GSK)^4^. Cynomolgus macaques are known to be suitable models for studying vaccine efficacy after SARS-CoV-2 infection because they develop pathological features of viral pneumonia and mild disease similar to natural human infections, with a high viral load in both the upper and lower respiratory tract^29^.

Both binding and neutralising antibodies are thought to be potential correlates of protection against COVID-19^30^. Indeed, a correlation has been demonstrated between vaccine-induced neutralising antibody titres and protective efficacy for several vaccines including Moderna’s mRNA-1273^31–33^. Some studies on transfusion of plasma from animal models or convalescent patients also confirmed the critical role of humoral immunity in protection against SARS-CoV-2^34, 35^. In the present study, vaccination with one dose of PHH-1V was enough to induce an early IgG antibody response against RBD in cynomolgus macaques. Likewise, two doses of PHH-1V induced higher RBD-specific IgG antibody levels one-week post-boost that were sustained until the end of the study (D42). PHH-1V vaccination also induced moderate RBD-specific IgA antibody in BAL on D42 (6 dpc), and there was a trend towards higher specific IgA antibody levels on D28 (pre-challenge) in PHH-1V-vaccinated animals. It has been previously reported that IgA antibodies could be involved in early virus neutralisation in the lower respiratory tract during SARS-CoV-2 infections, thus contributing to control viral replication at the mucosal surface^36^. The increase of IgA antibody levels observed on D42 compared to levels on D28 suggest a specific IgA response induced by SARS-CoV-2 infection, which was previously associated with a reduction in viremia after challenge^37^. Furthermore, PHH-1V vaccine has been shown to induce a potent neutralising activity 15 days post-boost against HIV-reporter pseudoviruses expressing SARS-CoV-2 spike from Beta, Delta and Omicron (BA.1, BA.4 and BA.5) variants, and against replicative SARS-CoV-2 D614G and VOCs Alpha, Gamma and Delta. Remarkably, our data confirms the cross-reactivity of the PHH-1V vaccine against Omicron (BA.5), the circulating variant at the time of writing this work.

Although there is no clear evidence linking CD4^+^ T-cell response with protection against SARS-CoV-2, some studies have shown a correlation of CD4^+^ T-cell responses with the magnitude of specific IgG and IgA against spike and live virus neutralising antibodies, which have been identified as correlates of protection^38, 39^. Nevertheless, a recent study of CD8^+^ cell depletion in macaques demonstrated the relevance of the CD8^+^ T-cell response in mediating protection when humoral responses are suboptimal^34^. In our study, prime-boost vaccination with PHH-1V elicited a robust CD4^+^ and CD8^+^ T-cell response with secretion of IFNL, TNFL and/or IL-2 against Alpha and Beta variants. Most important, PHH-1V elicited IFNL-secreting cells against Omicron (BA.1) spike, which highlights the cross-reactivity of immune responses elicited by PHH-1V vaccine against the most present variant at the time of writing of conducting this experiment. While PHH-1V vaccination induced a Th1/Th2-balanced T-cell response against Alpha and Beta spike, with similar levels of IFNL- and IL-4-secreting cells, it induced a Th1-skewed response against Omicron (BA.1) spike with a low detection of IL-4-secreting cells. Overall, SARS-CoV-2-specific T-cell responses induced in convalescent patients were mainly skewed towards Th1 cells, with the production of different Th1-like cytokines^38–40^. Indeed, it has been suggested that this Th1 cell-biased phenotype is associated with less severe disease^41^.

Efficacy of PHH-1V vaccine was assessed in cynomolgus macaques by the analysis of gRNA, sgRNA and infectious viral load in different tissues after SARS-CoV-2 infection. Two doses of PHH-1V significantly reduced SARS-CoV-2 gRNA in lungs and BAL compared to PBS mock-vaccinated animals. No SARS-CoV-2 gRNA was detected 6 dpi in lungs, trachea, oropharynx and nasal turbinates from most of PHH-1V vaccinated animals. It was also not detected in the BAL from three animals 6 dpi. These results are in line with the good levels of IgA antibodies detected in BALs before challenge. Likewise, animals vaccinated with PHH-1V cleared virus faster than control animals in nasopharyngeal swabs, nasal turbinate and trachea. Remarkably, infectious SARS-CoV-2 were undetectable after challenge in oropharyngeal and nasopharyngeal swabs (except for one animal at 1 dpc), BAL (except for one animal 1 at dpc) and lungs in PHH-1V vaccinated animals. Furthermore, sgRNA was also undetectable in BAL from PHH-1V-vaccinated animals, and less detected in nasopharyngeal swabs from vaccinated animals compared to non-vaccinated. Hence, PHH-1V not only prevents SARS-CoV-2 replication in the lower respiratory tract, but also decreases viral replication in the upper tract, which is especially important in reducing viral spread. This is of particular importance since protection in both the upper and lower respiratory tract is required to prevent transmission and disease in humans^42^. Non-clinical studies of adenovirus and mRNA vaccines showed persistent virus in nasal swabs despite preventing COVID-19^43^. For instance, studies in NHP with the Moderna’s mRNA-1273 or ChAdOx1 nCoV-19 vaccines demonstrated persistent viral RNA in nasal swabs of several animals^44, 45^. Although no infectious virus was detected in nasal swab samples of PHH-1V vaccinated animals from 2 dpc, low levels of viral RNA were still detected in three vaccinated animals at the end of the study. Further studies are required to elucidate whether our vaccine protects against SARS-CoV-2 transmission.

In line with no detectable viral load in the lungs of animals vaccinated with PHH-1V, we did not find bronchointerstitial inflammation and oedema in lung sections of vaccinated animals. Indeed, PHH-1V vaccine significantly reduced lung inflammation score compared to placebo immunisation. Similar results were reported in NHP studies with currently licensed vaccines^45, 46^. Of note, all non-vaccinated animals tested by IHC were positive in at least one lung section tested. Positive IHC results further support the evidence of SARS-CoV-2 replication in lungs of inoculated monkeys which was demonstrated by viral titration.

Additionally, PHH-1V is safe and well-tolerated since rectal temperature and body weight were stable throughout the study, and neither local reaction in the site of injection nor significant clinical signs were found. The changes in body temperature experienced by some animals during vaccination days can be explained by physiological temperature regulation of cynomolgus macaques over the day.

In summary, PHH-1V vaccine is safe, highly immunogenic against different VOCs including Omicron (BA.4 and BA.5), protects against SARS-CoV-2 replication in lower respiratory tract and controls viral infection in upper respiratory tract of cynomolgus macaques. Our data highlight the potential use of the PHH-1V vaccine in humans, which authorisation as a booster has been recently recommended by EMA’s human medicines committee (<u>CHMP</u>) in people aged 16 years and above who have been vaccinated with an mRNA COVID-19 vaccine .

## LIMITATIONS OF THE STUDY

Long-lasting protective efficacy were not assessed 6 months after SARS-CoV-2 challenge in this study, although long-term immunity is being analysed in phase IIb (NCT05305573) and phase III (NCT05303402) clinical trials. Additionally, protective efficacy after a prime-boost vaccination with PHH-1V has not been showed in this NHP model against SARS-CoV-2 beta, the variant from which the PHH-1V vaccine antigen is derived.

## ACKNOWLEDGMENTS

Anna Moya and Mireia Muntada for the ELISA analysis; Clara Panosa and Ester Puigvert for her assistance in the production of the vaccine antigen; Glòria Pujol and Eduard Fossas for their assistance in review of the manuscript; and Adrián Lázaro-Frías from Evidenze Health España S.L. for providing medical writing support during the preparation of this paper funded by Hipra Scientific, S.L.U. This project was partially funded by the Centre for the Development of Industrial Technology (CDTI, IDI20210115), a public organisation answering to the Spanish Ministry of Science and Innovation.

## AUTHOR CONTRIBUTIONS

**Conceptualisation, Investigation, Methodology, Resources, Formal Analysis and Data Curation:** Antoni Prenafeta, Gregori Bech-Sàbat, Alexandra Moros, Antonio Barreiro, Alex Fernández, Manuel Cañete, Mercè Roca, Luis González, Carme Garriga, Ricard March, Teresa Prat and Laura Ferrer; **Writing Original Draft, Writing – Review & Editing, Visualisation and Supervision:** Antoni Prenafeta and Alexandra Moros; **Investigation:** Antoni Prenafeta, Gregori Bech-Sàbat, Alexandra Moros, Antonio Barreiro, Alex Fernández, Manuel Cañete, Mercè Roca, Luis González, Carme Garriga, Edwards Pradenas, Silvia Marfil, Julià Blanco, Paula Cebollada, Marta Sisteré, Andreas Meyerhans, Joachim Confais, Marion Toussenot, Hugues Contamin, Andrés Pizzorno, Manuel Rosa-Calatrava, Cristina Lorca, Joaquim Segalés, Teresa Prat, Ricard March and Laura Ferrer; **Resources:** Edwards Pradenas, Silvia Marfil, Julià Blanco, Paula Cebollada, Marta Sisteré, Andreas Meyerhans, Joachim Confais, Marion Toussenot, Hugues Contamin, Andrés Pizzorno, Manuel Rosa-Calatrava, Cristina Lorca and Joaquim Segalés; **Writing – Review & Editing:** Antoni Prenafeta and Alexandra Moros; **Supervision:** Teresa Prat, Ricard March, and Laura Ferrer; **Project Administration:** Laura Ferrer; **Conceptualisation:** Antoni Prenafeta, Gregori Bech-Sàbat and Carme Garriga.

## STAR METHODS

### KEY RESOURCE DATA

Reagent and resource table is attached as a separate document.

### RESOURCE AVAILABILITY

#### Lead contact

Requests for further information or data should be directed to and will be fulfilled by the lead contacts, Antoni Prenafeta and Gregori Bech-Sabat.

#### Materials availability

Project-related biological samples are not available since they may be required by Regulatory Agencies or by HIPRA during the clinical development of the vaccine.

#### Data and code availability

- Data reported in this study cannot be deposited in a public repository because the vaccine is under clinical evaluation. Upon request, and subject to review, the lead contact will provide the data that support the reported findings.
- This paper does not report original code.
- Any additional information required to reanalyse the data reported in this paper is available from the lead contact upon request.

### EXPERIMENTAL MODEL AND SUBJECT DETAILS

#### Ethic statements

All procedures were conducted in accordance with the European Union Guidelines for Animal Welfare (Directive 2010/63/EU) and approved by the Ethics Committee of HIPRA Scientific S.L.U., the Animal Welfare Body of Cynbiose and the Ethics Committee of VetAgro-Sup (1 avenue Bourgelat, 69 280 Marcy l’Étoile, France) (approval number 2103; MESR number: 2020122910523878_v2). The animal facility is accredited by the Association for Assessment and Accreditation of Laboratory Animal Care (AAALAC) since 2015, an accreditation renewed in 2018 and 2021. This study was performed in compliance with Cynbiose standard operating procedures and according to the company quality management system.

#### Animal model

Twelve naïve cynomolgus macaques (*Macaca fascicularis*, 26 to 30 months old, 2.4 to 2.85 kg) were imported from Vietnam by Bioprim and were allocated to 2 groups (n=6; 3F + 3M) to assess immunogenicity, efficacy, and safety of PHH-1V. Animals were housed within the area dedicated to Cynbiose, in the animal facility owned by the veterinary school VetAgro Sup and managed by the Company Biovivo. Before experimental procedures, there was an acclimation period of at least 4 weeks to ensure compliance during manipulations. Litter (poplar wood shavings) maintenance was performed daily, and litter change and cleaning of the animal room were done at least once a month.

All animals were at first housed as isosexual social groups in a BSL-1 environment (irrespective of their treatment group) and were taken to a BSL-3 environment one day prior to their infection. From this point onward, animals were housed as separate isosexual treatment groups.

Regardless of the housing conditions, the environmental parameters were as follows: 12 hours light/dark cycle (temperature: 22 °C ± 2 °C, humidity: 30%–70%, at least 10 air changes per hour with no recirculation). Specific Primate Diet (Ssniff ref: V3944-000) was provided daily in appropriate amount (100 g for animals under 5 kg), as well as fresh fruits and vegetables. Tap water was available *ad libitum* to each animal via a water bottle with a sipper tube. Environmental enrichment was provided in the form of treats scattered in the litter daily and continuous access to toys, changed every week. Cells

Vero E6 cells (ATCC CRL-1586) were used for SARS-CoV-2 amplification and quantification of infectious viral load by TCID_50_. Cells were maintained in Dulbecco’s Modified Eagle’s Medium high glucose (DMEM, Lonza) with 4.5 g/l glucose, 5% Foetal Bovine Serum (FBS), 2mM of L-glutamine and 100 U/ml penicillin-streptomycin at 37 °C and 5% CO_2_.

HEK293T cells overexpressing WT human ACE-2 (Integral Molecular, USA) were used as target in pseudovirus-based neutralisation assay. Cells were maintained in T75 flasks with Dulbecco’s Modified Eagle’s Medium (DMEM) supplemented with 10% FBS and 1 μg/ml of puromycin (Thermo Fisher Scientific, USA).

Expi293F cells (Thermo Fisher Scientific) are a HEK293 cell derivative adapted for suspension culture that were used for SARS-CoV-2 pseudovirus production. Cells were maintained under continuous shaking in Erlenmeyer flasks following manufacturer’s guidelines.

#### Virus

SARS-CoV-2 B.1.160 variant (hCoV-19/Europe/France/ARA-CHUST-GIMAP-5/2020) encoding the spike mutation D614G was amplified on Vero E6 cells (batch number: D614G-P6-19/10/2021 Cynbiose) and quantified at 10^6.20^ TCID_50_/ml or 5.02×10^5^ PFU/ml. The genomic sequence of D614G strain is available on the EpiCoV GISAID platform (reference: EPI_ISL_1785075). B.1.1.7 variant (Alpha variant; hCoV-19/Europe/France/ARA-CHUST-GIMAP-10/2020) was amplified on Vero E6 cells (batch number: UK B117 P3 290321) and quantified at 10^6.53^ TCID_50_/ml. The genomic sequence is available on the EpiCoV GISAID platform (reference: EPI_ISL_1785073). P1 variant (Gamma variant; hCoV-19/French Guiana/IPP03772i/2021) was amplified on Vero E6 cells (batch number: BRS3772-130421P4) and quantified at 10^5.97^ TCID_50_/ml. The genomic sequence is also available on the EpiCoV GISAID platform (reference: EPI_ISL_1096262). Finally, B.1.617.2 strain (Delta variant) was amplified on Vero E6 cells (batch number IND-130721) and quantified at 10^6.97^ TCID_50_/ml. The sequence (available on demand) will be deposited on the EpiCoV GISAID platform.

### METHOD DETAILS

#### PHH-1V vaccine

The purified RBD fusion heterodimer was adjuvanted with an oil-in-water emulsion based on squalene. The PHH-1V vaccine was tested at 40 µg of RBD fusion heterodimer/dose for the safety and immunogenicity assays in cynomolgus macaques (*Macaca fascicularis*). The placebo vaccines consisted of phosphate buffered saline (PBS; Gibco). The staff conducting the *in vivo* and viral experiments was blinded to the treatment received by the animals. Furthermore, the staff receiving, preparing and administering the vaccine to the animals did not take part in the protocol afterwards.

#### Animal vaccination and SARS-CoV-2 challenge

Twelve animals (six per group) were intramuscularly immunised twice with 0.5 ml of PHH-1V vaccine or PBS (placebo) on D0 and D21 (prime-boost schedule). First administration was applied in right hindlimb and second in left hindlimb using sterile 1 ml syringes (Terumo® Syringe) and 25G needles (BD Microlance). Local reactions at the intramuscular injection sites were monitored the day of injection and daily until 5 dpi. Next, all animals were challenged on day 36 with 2 x 10^6^ PFU of SARS-CoV-2 (hCoV-19/Europe/France/ARA-CHUST-GIMAP-05/2020 [B.1.160 strain]) by intranasal (about 500 μl/nostril) and intratracheal route (4 ml) with a Penn-Century microsprayer connected to a safety syringe (Luer Lock).

Animals were monitored for clinical signs daily and for mortality twice daily. Bodyweight was recorded weekly and three times a week after infection, and food consumption was assessed daily for all animals housed in the same enclosure. Body temperature was monitored with a rectal thermometer around the administration (prime: from D-2 to D3; boost: from D19 to D24) and infection days (from D34 to D42). Peripheral blood was collected on days 0, 21, 28 and 36 before infection; and 1, 2, 3 and 6 dpc, by venepuncture from femoral veins with a Vacutainer® kit (Becton Dickinson). Blood samples were then used for serological analysis, PBMC preparation, and routine haematological and biochemical tests. BAL were collected 0, 2, 4 and 6 dpc for humoral immunity analysis and viral load assessment. Oropharyngeal, nasopharyngeal, and rectal swabs were also collected daily after infection for viral load measurement. Finally, animals were euthanised 6 dpc to viral load measurement and pathological examination of the lungs and other extrapulmonary tissues.

#### Haematological and biochemical evaluation

Blood was sampled into K3EDTA tubes (Vacuette®, Greiner Bio-One) for haematological analysis using a ProCyte Dx (Idexx, USA), and into Lithium Heparin tubes (BD Vacutainer^®^) for biochemical analysis with a Catalyst One (Idexx, USA) at different time points. Haematological analysis included red blood cell (RBC) count, white blood cell (WBC) count [including neutrophil (N), eosinophil (E), basophil (B), lymphocyte (L) and monocyte (M) count], haemoglobin concentration (HGB), haematocrit (HTC), mean corpuscular volume (MCV), mean corpuscular haemoglobin concentration (MCHC), mean corpuscular haemoglobin (MCH), platelet count (PLT), plateletcrit value (PCT), platelet large cell ratio (P-LCR), mean platelet volume (MPV), and reticulocytes (RET). Biochemical analysis included albumin (ALB), globulin (GLOB), alkaline phosphatase (ALT), alanine aminotransferase (ALT), calcium (Ca), cholesterol (CHOL), creatine kinase (CK), creatinine (CREA), gamma glutamyl transferase (GGT), glucose (GLU), inorganic phosphates (PHOS), total bilirubin (TBIL), total protein (TP), triglycerides (TRIG), urea (UREA) and blood electrolytes [including sodium (Na), chloride (Cl) and potassium (K)].

#### Sample collection for SARS-CoV-2 quantification

##### Bronchoalveolar lavage (BAL) collection

BAL were obtained by infusion of 5 ml/kg of a solution of isotonic sodium chloride (NaCl 0.9%; B. Braun) into the trachea, kept at approximately 37 °C until use to avoid thermal shock of the mucosa. Supernatant and cell pellet from BAL samples were separated by centrifugation at 2500 rpm for 5 minutes at 4 °C. Cell pellets were stored at 5 °C ± 3 °C in RNA*later* (Thermo Fisher Scientific) for measurement of RNA viral copies by genomic RT-qPCR, or in DPBS without Ca and Mg (Sigma Life Science) for detection of infectious virus. Supernatants were immersed in a hot water bath (56 °C) for 30 minutes to inactivate the virus before freezing at -20 °C, and were used for IgA and subgenomic RT-qPCR determinations.

##### Nasopharyngeal, oropharyngeal and rectal swab collection

Animal were sedated and nasopharyngeal (right and left nostril), oropharyngeal and rectal samples were collected with specific swabs (Puritan®, Hydraflock micro ultrafine swabs) and stored at 5 °C in RNA*later* until processing.

##### Organ collection

Pieces of organs were collected and then placed in 1.2 ml cryotubes full of *RNAlater* for viral load analysis by RT-qPCR or placed in dry tubes without storage medium for TCID_50_ assays (pieces of organs were weighed before packaging for TCID_50_ analysis). Next, organs were washed in PBS and crushed with forceps and scalpel before disruption by a Tissue Lyser LT (Qiagen) during 10 minutes in presence of two tungsten carbide beads (Qiagen). Samples intended for TCID_50_ were grinded in cold PBS and samples intended for RT-qPCR in RLT lysis buffer from the RNeasy Mini Kit (Qiagen). All samples were then stored at 5 °C.

#### Quantification of genomic RNA by RT-qPCR

Relative quantification of viral genome was performed by one-step real-time quantitative reverse transcriptase and polymerase chain reaction (RT-qPCR) from total RNA extracted from samples preserved in RNA*later* using the RNeasy Mini kit (QIAGEN). Real-time one-step RT-qPCR was performed using the EXPRESS One-Step Superscript™ qRT-PCR Kit (Invitrogen), as previously described^47^. Briefly, we prepared a 20 µl reaction volume containing 10 µl of Express qPCR supermix at 2X, 1 µl of forward primer at 10 mM, 1 µl of reverse primer at 10 mM, 0.5 µl of probe at 10 mM, 3.1 µl of PCR-water (QIAGEN), 0.4 µl of Rox dye at 25 mM, and 2 µl of RNA template. Primers and probe sets were designed by the School of Public Health (University of Hong Kong)^48^ and synthetised by Eurogentec. The set is specific to the non-structural protein 14 (ORF1b-nsp14) gene of SARS-CoV-2 (forward primer HKU-ORF1b-nsp14F: 50-TGGGGYTTTACRGGTAACCT-30; reverse primer HKUORF1b-nsp14R: 50-AACRCGCTTAACAAAGCACTC-30; probe HKU-ORF1b-nsp141P: 50-FAM-TAGTTGTGATGCWATCATGAC TAG-TAMRA-30). Gene expression assay of the simian GAPDH (housekeeping gene) was also performed as endogenous control.

Thermal cycling and data collection of RT-qPCR were performed in QuantStudio™ 5 Real-Time PCR System (Applied Biosystems) in MicroAmp™ Fast Optical 96-well reaction plates (Applied Biosystems). Cycling conditions were as follows: reverse transcription at 50 °C for 15 minutes, followed by initial polymerase activation at 95 °C for 2 minutes^49^, and then 40 cycles of denaturation at 95 °C for 15 seconds and annealing/extension at 60 °C for 1 minute. Analysis was done with the QuantStudio™ Design and Analysis Software v1.5.1. Genomic RNA was expressed as log_10_ (nsp14 copies/mL) in BAL and swabs or as log_10_ (nsp14 copies/GAPDH copies) in tissues.

#### Quantification of subgenomic RNA by RT-qPCR

Viral RNA from BAL was extracted using the IndiMag pathogen kit (Indical Biosciences, Germany) on a Biosprint 96 workstation (Qiagen, Germany), according to the manufacturer’s instructions. Subgenomic RNA detection by RT–qPCR from BAL and NPS samples was assessed using the primers: forward 5′-CGATCTCTTGTAGATCTGTTCTC-3′, reverse 5′-ATATTGCAGCAGTACGCACACA-3′ and the probe 5′-FAM-ACACTAGCCATCCTTACTGCGCTTCG-TAMRA-3′. The concentration of each of the primers was 400 nM and 200 nM for the probe as previously described^50^, with minor modifications from Wölfel et al. to adapt it to AgPath-ID One-Step RT–PCR Kit^51^. Thermal cycling was performed at 50 °C for 10 minutes for reverse transcription, followed by 95 °C for 10 minutes and then 45 cycles of 95 °C for 15 seconds, 56 °C for 30 seconds by using a 7500 Fast Real-Time PCR System (Applied Biosystems, Life Technologies, USA). Samples with a Cq value ≤ 40 were considered positive for SARS-CoV-2.

#### Quantification of infectious viral load by TCID_50_

TCID_50_ assays were performed with confluent Vero E6 cells in 96-well microplates (Thermo Fisher Scientific) using 50 μl of ten-fold serial dilutions (10^-1^ to 10^-6^) of virus samples in quadruplicate. Plates were then incubated for 4 days at 37 °C and 5% CO_2_ atmosphere and cytopathic effects were observed through an optical microscope (Nikon, Eclipse Ti). The total number of positive and negative wells for each dilution was recorded and TCID_50_ was calculated using the Reed and Muench method^49^. Viral load was expressed as log_10_ TCID_50_/ml in oropharyngeal and nasopharyngeal swabs and BAL, and log_10_ TCID_50_/g in lungs and other tissues.

#### Histopathology examination

Organ samples from all pulmonary lobes, the upper respiratory tract (trachea, oropharyngeal and nasal mucosa), lymph nodes, kidney, heart, liver, spleen, stomach, and colon were collected for histopathology and preserved in 4% PFA solution (Sigma). Samples were embedded into paraffin blocks and then sectioned by microtome at approximately 4 μm. Sections were then stained with haematoxylin and eosin.

Individual microscopic observations were recorded including a severity grade indicating extension and/or severity according to a 5-tier scoring system (minimal, mild, moderate, marked, and severe). Overall, a score ≤ 3 indicates minimal microscopic changes, a score from 3 to 10 indicates mild to moderate inflammation and a score ≥10 indicates extensive inflammation. Selected microphotographs were captured to illustrate microscopic observations in the lung.

#### Immunohistochemistry analysis

Lung tissue embedded in paraffin were cut at 4 µm and a specific immunohistochemistry protocol to detect the nucleocapsid protein of SARS-CoV-2 was performed. A rabbit monoclonal antibody targeting the nucleocapsid protein of SARS-CoV-2 (40143-R019, Sino Biological Inc., China; dilution 1:1000) was used to detect the presence of SARS-CoV-2 antigen, as previously described^50^. The amount of viral antigen in tissue samples was semi-quantitatively scored per section: lack of antigen detection (score 0), very low (score 1, 1-2 small clusters of positive cells), low (score 2, 3-5 clusters of positive cells in multifocal distribution), moderate (>5 clusters of positive cells in multifocal distribution), and high (score 4, multifocal to diffuse distribution of positive cells). Selected sections were captured to illustrate microscopic observations in the lung.

#### Analysis of SARS-CoV-2-specific IgG antibodies by ELISA

Serum binding antibodies against SARS-CoV-2 RBD were determined by ELISA. MaxiSorp plates (Nunc) were coated with 100 ng/well RBD protein (Sino Biologicals) and blocked with 5% non-fat dry milk (Difco, BD) in PBS. Wells were incubated with serial dilutions of the serum samples and the bound total IgG specific antibodies were detected by goat anti-monkey IgG (H+L) secondary antibody, HRP (Thermo Fisher). Finally, wells were incubated with K-Blue advanced substrate (Neogen) and the absorbance at 450 nm was measured using a microplate reader (Versamax, Molecular Devices). The mean value of the absorbance was calculated for each dilution of the serum sample run in duplicate. The end-point titre of RBD-specific total IgG binding antibodies was established as the reciprocal of the last serum dilution yielding 3 times the mean optical density of the negative control of the technique (wells without serum added).

#### Analysis of SARS-CoV-2-specific IgA antibodies by ELISA

IgA antibodies against SARS-CoV-2 RBD were determined by ELISA. MaxiSorp plates (Nunc) were coated with 200 ng/well RBD protein (Sino Biologicals) and blocked with StabilCoat™ Immunoassay Stabilizer (Surmodics). Wells were incubated with serial dilutions of the BAL samples and the bound total IgA specific antibodies were detected by goat anti-human IgA biotinylated (Mabtech) with cynomolgus macaque cross-reactivity. After washes, streptavidin-HRP (Mabtech) was employed for finalising IgA detection. Finally, wells were incubated with K-Blue advanced substrate (Nirco) and the absorbance at 450 nm was measured using a microplate reader (VersaMax, Molecular Devices). The mean value of the absorbance was calculated for each dilution of the BAL sample run in duplicate. The end-point titre of RBD-specific total IgA binding antibodies was established as the reciprocal of the last BAL sample dilution yielding 3 times the mean optical density of the negative control of the technique (wells without sample added).

#### Pseudovirus neutralisation assay

Neutralising antibodies in serum against SARS-CoV-2 Beta, Delta and Omicron (BA.1) variants were determined by a pseudoviruses-based neutralisation assay (PBNA) at IrsiCaixa AIDS Research Institute (Barcelona, Spain) using an HIV reporter pseudovirus that expresses the S protein of SARS-CoV-2 and luciferase. Pseudoviruses were generated as previously described^52^. For the neutralisation assay, 200 TCID_50_ of pseudovirus supernatant was preincubated with serial dilutions of the heat-inactivated serum samples for 1 hour at 37 °C and then added onto ACE2 overexpressing HEK293T cells. After 48 hours, cells were lysed with britelite plus luciferase reagent (PerkinElmer). Luminescence was measured for 0.2 seconds with an EnSight multimode plate reader (PerkinElmer). The neutralisation capacity of the serum samples was calculated by comparing the experimental RLU calculated from infected cells treated with each serum to the max RLUs (maximal infectivity calculated from untreated infected cells) and min RLUs (minimal infectivity calculated from uninfected cells) and expressed as the neutralisation percentage: Neutralisation (%) = (RLUmax–RLUexperimental)/(RLUmax–RLUmin) * 100. IC_50_ were calculated by plotting and fitting neutralisation values and the log of plasma dilution to a 4-parameters equation in Prism 9.0.2 (GraphPad Software, USA).

#### Microneutralisation assay

Neutralising antibodies against SARS-CoV-2 D614G, Alpha, Gamma and Delta variants were measured using a microneutralisation test (MNT) assay in the BSL-3 facility at VirNext (Lyon, France). Serum samples were two-fold serially diluted in DMEM-2% FBS medium and were incubated at a 1:1 ratio with 100 TCID_50_ of SARS-CoV-2 variants in 96-well microplates for 1 hour at 37 °C. Then, mixtures of serum samples and SARS-CoV-2 variants were added in duplicate to Vero E6 confluent monolayers seeded in 96-well microplates and were incubated at 37 °C in a 5% CO_2_ incubator for 4 days. The cytopathic effect was observed using an inverted microscope at 72- and 96-hours post-infection and the titre was calculated as the inverse of the last serum dilution producing no cytopathic effect on the cells.

#### IFNL and IL-4 ELISpot assays

The NHP IFNL and IL-4 ELISpot Plus kits were used following manufacturer’s instructions (Mabtech). In brief, a total of 2.5 x 10^5^ (for the IFNL assay) or 4 x 10^5^ (for the IL-4 assay) per well were plated in duplicates and *ex vivo* stimulated with a 1:1 mixture of RBD-overlapping peptides from the SARS-CoV-2 B.1.1.7 (Alpha) and B.1.351 (Beta) lineages, or BA.1 (Omicron) (JPT peptide technologies) (1 µg/ml per peptide final concentration). As controls, PBMCs were incubated with complete RPMI (negative control), or with 2.5 ng/ml PMA plus 250 ng/ml ionomycin (positive control; both Sigma). After 18-20 hours incubation (for the IFNL) and 48 hours incubation (for the IL-4), plates were developed using kit reagents. Spots were counted under a dissection microscope (Leica GZ6). Frequencies of IFNL- and IL-4-producing cells were calculated by subtracting the mean of spots counted in the RBD peptide-stimulated wells from the mean of spots counted in the negative control wells. Data was expressed as number of RBD-specific IFNL-secreting cells per 1 × 10^6^ PBMCs or RBD-specific IL-4-secreting cells per 1 x 10^6^ PBMCs.

#### Intracellular cytokine staining (ICS)

Isolated cynomolgus macaques PBMCs in RPMI (Gibco) supplemented with 10% FBS, 1% penicillin/streptomycin, 0.05 mM β-mercaptoethanol and 1 mM sodium pyruvate (cRPMI) were seeded into a 96-well round bottomed plate (1×10^6^ macaque PBMCs in 100 μl volume per well) and stimulated under three different conditions: (i) a 1:1 mixture of the peptide libraries (PepMix™) (JPT peptide technologies) from SARS-CoV-2 lineages B.1.1.7 (Alpha) and B.1.351 (Beta) covering the RBD of the Spike protein, (ii) cRPMI (negative control) or (iii) PMA + ionomycin (positive control). Incubation was for 5 hours at 37 °C and 5% CO_2_. Brefeldin A (10 μg/ml; BFA, Sigma) was added for the last 3 hours to block cytokine secretion. Final concentrations of individual peptides of the RBD peptide pools were 1 μg/ml. PMA (Sigma) and ionomycin (Sigma) were used in a final concentration of 2.5 ng/ml and 250 ng/ml, respectively.

For cell and cytokine characterisation by flow cytometry, cells were first labelled with Zombie NIR fixable viability stain (BioLegend) in PBS for 15 minutes on ice, and then stained with fluorescence-labelled monoclonal antibodies CD3-AF700 (clone SP34-2, BD Bioscience), CD4-BV650 (clone L200, BD Bioscience), CD8-Pacific Blue (clone RPA-T8, BD Bioscience) and CD69-PeCy7 (clone FN50, eBioscience) for 20 minutes on ice in FACS buffer (PBS 5% FCS, 0.5% BSA, 0.07% NaN3). Cells were subsequently fixed in 2 % formaldehyde for 20 minutes on ice and stained for intracellular cytokines for 20 minutes on ice with IFNL-APC (clone B27, BD Bioscience), IL-2-PE (clone MQ1-17H12, BD Bioscience), TNF-AF488 (clone mAb11, BioLegend), IL-4 BV421 (clone MP4-25D2, BD Bioscience) antibodies diluted in Perm/Wash buffer (PBS 1% FCS, NaN3 0.1%, Saponin 0.1%). Thereafter, a second intracellular staining step of 10 minutes on ice was performed using the Streptavidin PE-Cy7 conjugate (eBioscience, Thermofisher Scientific).

Fluorescence-labelled cells were analysed on an Aurora (Cytek) flow cytometer followed by data analysis using FlowJo 10 software (Tre Star Inc). The gating strategy followed in the analysis is showed in **Figure S1**. RBD-specific cytokine-positive responses were obtained by subtracting background cytokine expression from the negative control stimulations (cRPMI only). To calculate the Th1 response in CD4^+^ and CD8^+^, the Boolean tool of the FlowJo software was used.

#### Correlation studies

To test the association between the immunogenicity and efficacy parameters, Kendall’s Rank correlation tests have been employed. This method is recommended for ordinal and continuous variables that are not strictly normally distributed and/or are not linearly (but monotonically) correlated. Immunological parameters chosen for this analysis were the log_10_ IC_50_ antibody titres against SARS-CoV-2 D614G and beta variant at day 36 from VNA and PBNA, respectively; and IFN-γ-secreting cell levels after alpha-beta and omicron stimulation. Efficacy parameters chosen for this analysis were the log_10_ TCID_50_/ml titres of the nasopharyngeal and oropharyngeal swabs, BAL and different lung sections. To calculate the Kendall’s τ_B_ correlation coefficient and its 95% confidence interval the cor.test() function in R with the argument method = “kendall” and the KendallTauB() function of the DescTools R package have been used.

### QUANTIFICATION AND STATISTICAL ANALYSIS

Statistical analyses and plots were generated using R (version 4.0.5) or GraphPad Prism (version 9). Unless otherwise specified, all plots depict individual data points for each animal, along with the sample mean and standard deviation. When required, data was either log_10_- or arcsine-transformed (i.e., log-normal and percentage variables, respectively). The exact number (n) used in each experiment is indicated in the caption below each figure.

When testing the effect of one or two factors, linear models were generally employed. For models involving independent observations, the generalised least squares approximation (GLS implementation in the R package nlme) was used to accommodate potential heteroskedasticity. Conversely, for models involving repeated measures, linear mixed effects models were fitted using the lme implementation in the R package nlme. Unless otherwise specified, time, group and their interaction were included in the models as fixed effects, and the experimental subject was considered a random factor. The corresponding random intercept models were fitted to the data using restricted maximum likelihood. Correlation between longitudinal observations as well as heteroskedasticity were included in the models when required with appropriate variance-covariance structures.

Assumptions were tested graphically (using quantile-quantile and residual plots) for both modelling approaches, and model selection was based on likelihood ratio tests or a priori assumptions. The corresponding estimated marginal means were calculated and compared using the R package emmeans.

On the other hand, data violating the assumption of normality were analysed using a Welch’s permutation t-test or Mann-Whitney’s U test, segregating by timepoint if needed.

When comparisons against zero-variance groups (all observations having the same value) needed to be performed, one-sample tests were employed instead.

Statistically significant differences between groups are indicated with a line on top of each group: ** p<0.01; * p<0.05.

