## Supplementary material for "Preclinical evaluation of PHH-1V vaccine candidate against SARS-CoV-2 in non-human primates"

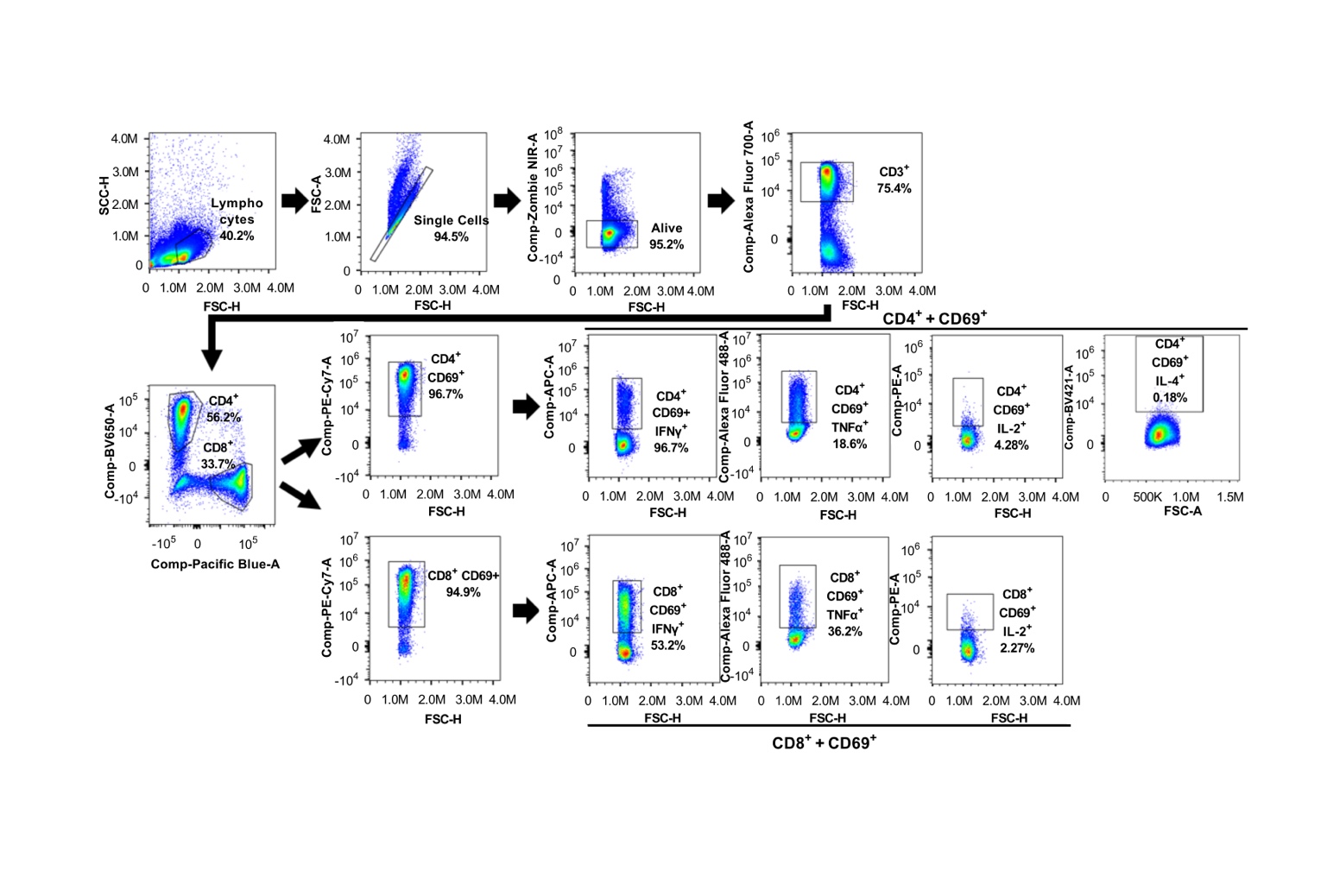

**Figure S1. Gating strategy to identify IFNγ, TNFα, IL-2 and IL-4-secreting CD4^+^ T cells and IFNγ, TNFα and IL-2-secreting CD8^+^ T cells from cynomolgus macaques splenocytes by flow cytometry.** Related to Figure 3. FSC-H/FSC-A was used to exclude doublets, and T cells (CD3^+^) within the alive cells (LIVE DEAD Far Red^-^) were gated. Then, IFNγ, TNFα, IL-2 and IL-4-producing T cells upon mock or RBD-stimulation were measured from CD4^+^ CD69^+^ and/or CD8^+^ CD69^+^ populations. Fluorescence Minus One (FMO) controls were used to verify flow cytometric data.

| **Nasopharyngeal swab** | | | | | | | | **BAL supernatant*** | | | |
| --- | --- | --- | --- | --- | --- | --- | --- | --- | --- | --- | --- |
| **Group** | **Pre-challenge** | **1 dpc** | **2 dpc** | **3 dpc** | **4 dpc** | **5 dpc** | **6 dpc** | **Pre-challenge** | **2 dpc** | **4 dpc** | **6 dpc** |
| control | undetected | 32.8 | undetected | undetected | undetected | undetected | undetected | undetected | 33.9 | 35.2 | undetected |
| control | undetected | 38.5 | 29.2 | undetected | undetected | undetected | undetected | undetected | 31.8 | undetected | undetected |
| control | undetected | 31.3 | 30.9 | undetected | undetected | 38.4 | undetected | undetected | 34.6 | undetected | undetected |
| control | undetected | undetected | undetected | 32.4 | undetected | 36.3 | undetected | undetected | 36.0 | undetected | undetected |
| control | undetected | undetected | 28.6 | 28.8 | 28.7 | 32.7 | undetected | undetected | 35.7 | undetected | undetected |
| control | undetected | undetected | undetected | undetected | 36.2 | undetected | undetected | undetected | undetected | undetected | undetected |
| PHH-1V | undetected | undetected | undetected | undetected | undetected | undetected | 37.6 | undetected | undetected | undetected | undetected |
| PHH-1V | undetected | undetected | undetected | undetected | undetected | 35.9 | undetected | undetected | undetected | undetected | undetected |
| PHH-1V | undetected | undetected | undetected | undetected | undetected | undetected | undetected | undetected | undetected | undetected | undetected |
| PHH-1V | undetected | undetected | undetected | undetected | undetected | undetected | undetected | undetected | undetected | undetected | undetected |
| PHH-1V | undetected | undetected | undetected | undetected | 39.4 | undetected | undetected | undetected | undetected | undetected | undetected |
| PHH-1V | undetected | undetected | undetected | undetected | undetected | undetected | undetected | undetected | undetected | undetected | undetected |

**Figure S2. Detection of subgenomic RNA in nasopharyngeal swabs and BAL of infected animals.** Related to Figure 4. Subgenomic viral RNA (sgRNA) was analysed by RT-qPCR in nasopharyngeal swabs and BAL from PHH-1V-vaccinated (n=6) and control (n=6) animals on D28 (pre-challenge) and at different times post-challenge. Cycle threshold (Ct) was showed for every animal at different times post-challenge. Data shaded in red and blue show Ct values for control animals and PHH-1V-vaccinated animals, respectively. Fisher's Exact tests were employed to compare the proportion of shedding animals in the two groups. An animal was considered as shedder if at least one positive sgRT-qPCR test was observed during the timecourse. Ct values >40 are considered negative. (*p<0.05).

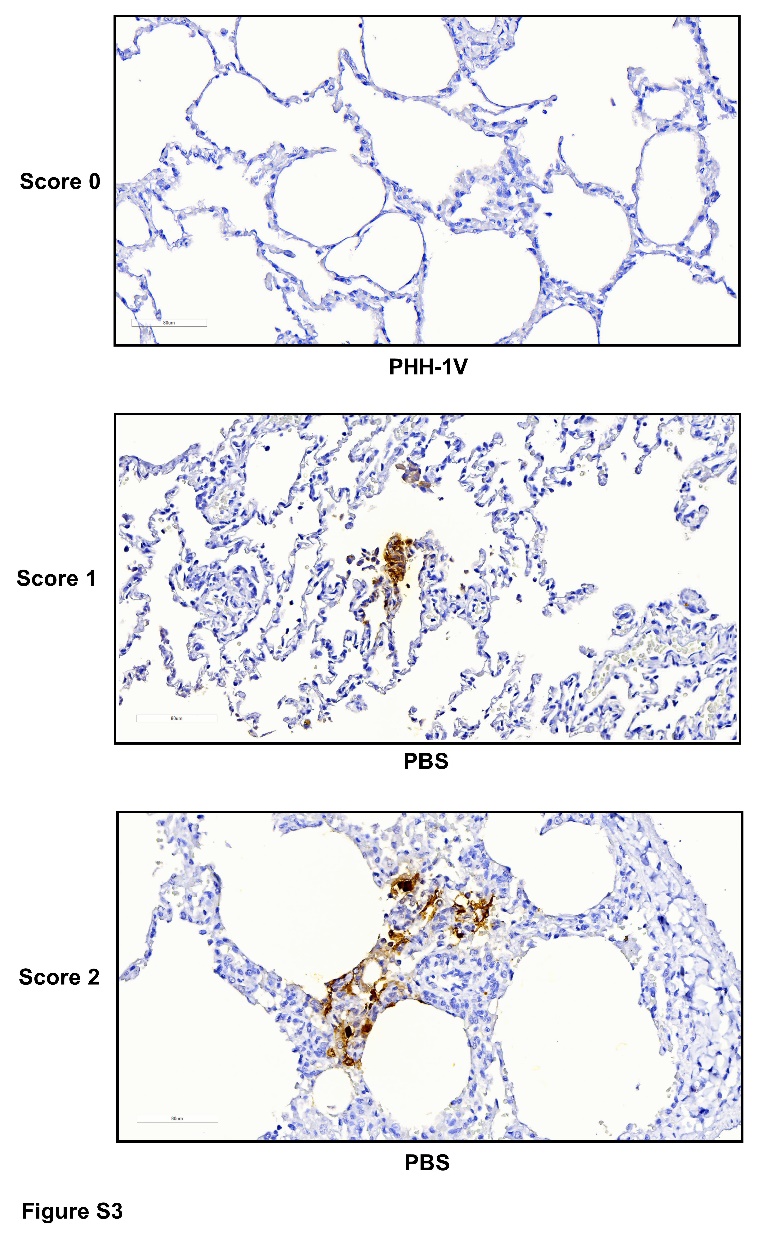
**Figure S3**. **Immunohistochemistry assessment in lung samples of infected animals.** Related to Figures 5 and 6. Immunohistochemistry to detect SARS-CoV-2 nucleoprotein in formalin-fixed, paraffin-embedded lung tissues from inoculated Cynomolgus macaques. All sections tested were from animals with a positive SARS-CoV-2 titre in the lung. The amount of viral antigen in tissue samples was semi-quantitatively scored per section: lack of antigen detection (score 0), very low (score 1, 1-2 small clusters of positive cells), low (score 2, 3-5 clusters of positive cells in multifocal distribution), moderate (>5 clusters of positive cells in multifocal distribution), and high (score 4, multifocal to diffuse distribution of positive cells). Selected sections were captured to illustrate microscopic observations in the lung.

| **Efficacy outcome** | **Immunogenicity parameters** | | | |
| --- | --- | --- | --- | --- |
| Viral load (TCID_50_) | nAb by PBNA  (IC_50_) | nAb by VNA  (end-point titre) | IFN-γ^+^ T-cells by ELISpot  (Alpha-Beta) | IFN-γ^+^ T-cells by ELISpot  (Omicron) |
| NPS | -0.39 (-0.733, -0.048)* | -0.49 (-0.822, -0.158)* | -0.36 (-0.854, 0.144) | -0.04 (-0.518, 0.445) |
| OPS | -0.26 (-0.662, 0.136) | -0.56 (-0.819, -0.297) | -0.61 (-0.933, -0.285) | -0.36 (-0.798, 0.085) |
| BAL | -0.37 (-0.728, -0.014)* | -0.64 (-0.827, -0.452) | -0.72 (-0.886, -0.560) | -0.62 (-0.883, -0.354) |
| RLCL | -0.46 (-0.894, -0.029) | -0.58 (-0.877, -0.290) | -0.52 (-0.891, -0.148) | -0.82 (-0.971, -0.663) |
| RLML | -0.37 (-0.868, 0.131) | -0.42 (-0.915, 0.068) | -0.49 (-0.796, -0.178)* | -0.65 (-1.000, -0.285) |
| LLCL | -0.29 (-0.622, 0.042) | -0.47 (-0.756, -0.191) | -0.71 (-0.939, -0.479) | -0.42 (-0.680, -0.165)* |
| LLCRLC | -0.21 (-0.634, 0.224) | -0.38 (-0.767, 0.007) | -0.71 (-0.959, -0.459) | -0.30 (-0.802, 0.213) |
| LLCRLCR | -0.58 (-0.850, -0.317) | -0.60 (-0.833, -0.363) | -0.56 (-0.907, -0.208) | -0.57 (-0.881, -0.262) |

**Figure S4. Summary table of correlation assessments.** Related to Figures 2, 3 and 5. A Kendall’s Rank correlation tests have been performed to assess the association between the immunogenicity parameters and efficacy outcome. The following efficacy outcomes were chosen for these analysis: viral load in nasopharyngeal swabs (NPS), oropharyngeal swabs (OPS), bronchoalveolar lavages (BAL), right lung caudal lobe (RLCL), right lung medial lobe (RLML), left lung caudal lobe (LLCL), left lung cranial lobe - caudal part (LLCRLC) and left lung cranial lobe - cranial part (LLCRLCR). Results were expressed as $\tau_{B}$ (95% CI). Data shaded in Gray show a statistically significant correlation between both parameters (p<0.05). *These correlation assessments were not significant at the 5% level but provided 95% confidence intervals completely below 0. Discrepancies between p-values and CIs are attributed to the presence of ties in the data.
